# Serotonin stimulated parathyroid hormone related protein induction in the mammary epithelia by transglutaminase-dependent serotonylation

**DOI:** 10.1101/2020.03.27.012195

**Authors:** Celeste M. Sheftel, Laura L. Hernandez

## Abstract

Mammary-derived serotonin [5HT] has been implicated in breast-to-bone communication during lactation by increasing parathyroid hormone related-peptide [PTHrP] in the mammary gland. It is well-established that PTHrP acts on the bone to liberate calcium for the milk during lactation; however, the mechanism of 5HT’s regulation of PTHrP has not been fully elucidated. Recently, serotonylation, has been shown to be involved in a variety of physiological processes. Therefore, we investigated whether serotonylation is involved in 5HT’s regulation of PTHrP in the mammary gland. Using lactogenic differentiated mouse mammary epithelial cells, we studied the effect of increased intracellular 5HT using the antidepressant, fluoxetine [FLX], or 5-hydroxytryptophan ([5HTP] 5HT precursor) with or without transglutaminase inhibition on PTHrP induction and activity and the potential serotonylation target protein, RhoA. Treatment with FLX or 5HTP significantly increased intracellular 5HT concentration and subsequently increased PTHrP gene expression which was reduced with transglutaminase inhibition. Further, we demonstrated that transglutaminase becomes more active with lactogenic differentiation and with 5HTP or FLX treatment. We examined RhoA, Rac1, and Rab4 as potential serotonylation target proteins and have concluded RhoA is likely a serotonylation target protein. Our data suggest that 5HT regulates PTHrP induction in part through the process of serotonylation during lactation.

## Introduction

Serotonin (5-hydroxytryptamine [5HT]), an established monoamine and neurotransmitter, is synthesized from L-tryptophan in a 2-step conversion requiring a hydroxylation step and a decarboxylation step. The hydroxylation is performed by the rate-limiting enzyme, tryptophan hydroxylase [TPH], producing 5-hydroxytryptophan [5HTP], which is decarboxylated by aromatic L-amino acid decarboxylase producing 5HT (1). 5HT is then released into the blood and stored by platelets, which lack the TPH1 enzyme. Degradation of 5HT occurs through oxidation by monoamine oxidase into 5-hydroxyindolacetic acid which is excreted in the urine. In humans and in mice, there are two TPH enzymes: TPH2 which converts L-tryptophan to 5HTP in the central nervous system and TPH1 which converts L-tryptophan to 5HTP in the periphery (2–4). 5HT and TPH1/2 are unable to cross the blood brain barrier, resulting in compartmentalization of 5HT (2,5). While 5HT is well-known as a central neurotransmitter altering behavior and mood, over 95% of 5HT is produced in the periphery in the gut enterochromaffin cells. During lactation, it has been has demonstrated that the mammary gland contributes approximately 50% of circulating 5HT (6,7).

Mammary-derived 5HT is important in regulating maternal calcium homeostasis and breast-to-bone communication via the synthesis and secretion of the parathyroid hormone related protein [PTHrP] in the mammary gland during lactation (8–10). PTHrP is secreted from the mammary epithelial cells and then acts on the bone to liberate calcium for milk synthesis (11–13). Since the fetus is not fully mineralized in utero, there is a disproportionate demand for calcium post-partum to synthesize milk. This results in utilization of the maternal bone to supply a large portion of the calcium for the baby through the milk, resulting in up to 10% maternal bone lost during the recommended 6 months of exclusive breastfeeding (14–17). Additional stressors on the mother, such as the use of antidepressants like selective serotonin reuptake inhibitors [SSRIs] have been demonstrated to result in a sustained reduction of trabecular bone density (18,19). Furthermore, our lab has shown *in vivo* exposure to the SSRI fluoxetine [FLX] leads to an increase in mammary gland 5HT content and consequently PTHrP (*Pthlh* gene) during lactation; further we determined maternal trabecular bone fails to restore post-weaning resulting in a sustained reduction in trabecular bone due to SSRI use (20).

SSRIs are the first-choice treatment for peripartum depression due to the low fetal teratogenicity and high patient compliance; often the benefits of the SSRI outweigh the potential negative effects of use (21–25). SSRIs exert their action by blocking the 5HT reuptake transporter, SERT, increasing the exposure of a neuron or tissue to 5HT. This can secondarily upregulate TPH1/2 increasing 5HT synthesis and decrease 5HT degradation (26). We have previously demonstrated an epigenetic-hedgehog link between 5HT and PTHrP involving DNA methylation (27). However, 5HT has no donatable methyl groups, suggesting a secondary mechanism or indirect regulation of DNA methylation.

Novel research in the past decade has elucidated a covalent post-translational modification involving 5HT, termed serotonylation (28). In this reaction, 5HT is transamidated onto a glutamine residue of a target protein via the calcium dependent enzyme, transglutaminase [TG] (29). In humans there are 9 TG genes, resulting in 8 active enzymes; however, TG2 [termed tissue transglutaminase] is ubiquitously expressed and is generally considered to be the most-likely TG enzyme involved in the transamidation of monoamines (29,30). Serotonylation was first described in platelet α-granule release through the activation of small GTPases (28). Since then, it has been implicated in vascular processes (i.e., smooth muscle contraction), pulmonary hypertension, glucose metabolism, dendritic spine remodeling in the brain, and most recently in histone modifications altering transcription (31–36). Small G-proteins such as Rho, Rab4, or Rac1 are common target proteins for serotonylation (28,34,37,38). In the Rho and Rab family of G-proteins, 5HT is transamidated onto the switch-2 domain, rendering them constitutively active until they are ubiquitinated for proteasomal degradation (30,37). TG2, the enzyme responsible for serotonylation, has been implicated in the promotion of growth, survival, and metastasis of breast cancer in mammary epithelial cells, however the molecular process of serotonylation has not been explored (39,40). Additionally, there has been no focus on the role of serotonylation during the normal lactogenic processes.

Here, we examined the role of TG2 in the mechanism involving 5HT’s regulation of PTHrP under lactogenic conditions. We hypothesized *in vitro* lactogenic differentiated mammary epithelial cells, using pharmacological manipulation to increase intracellular 5HT, will increase PTHrP via the molecular process, serotonylation. This regulation of PTHrP will be restored to control levels with TG inhibition. Further, we investigated the role of the potential serotonylation target, the small G protein RhoA.

## Results

### Transglutaminase protein and activity are observed in HC11 cells

We analyzed TG2 protein expression by immunoblot in both undifferentiated and lactogenically differentiated HC11 cells (Fig. 1A). TG2 is present in HC11 cells but is unchanged based on treatment with lactogenic hormones. This is further supported by *TG2* mRNA expression (Fig. 1B) which is not affected by lactogenic differentiation. In addition to the presence of TG2 expression at both the mRNA and protein levels in the HC11 cells, TG2 is active. Using a TG activity assay (Fig. 1C), we demonstrate that TG activity increases over time after treatment with lactogenic hormones. Together, these data demonstrate the enzyme required for serotonylation is present and active in mammary epithelial cells, suggesting the molecular process of serotonylation can occur in these cells.

**Figure 1:**
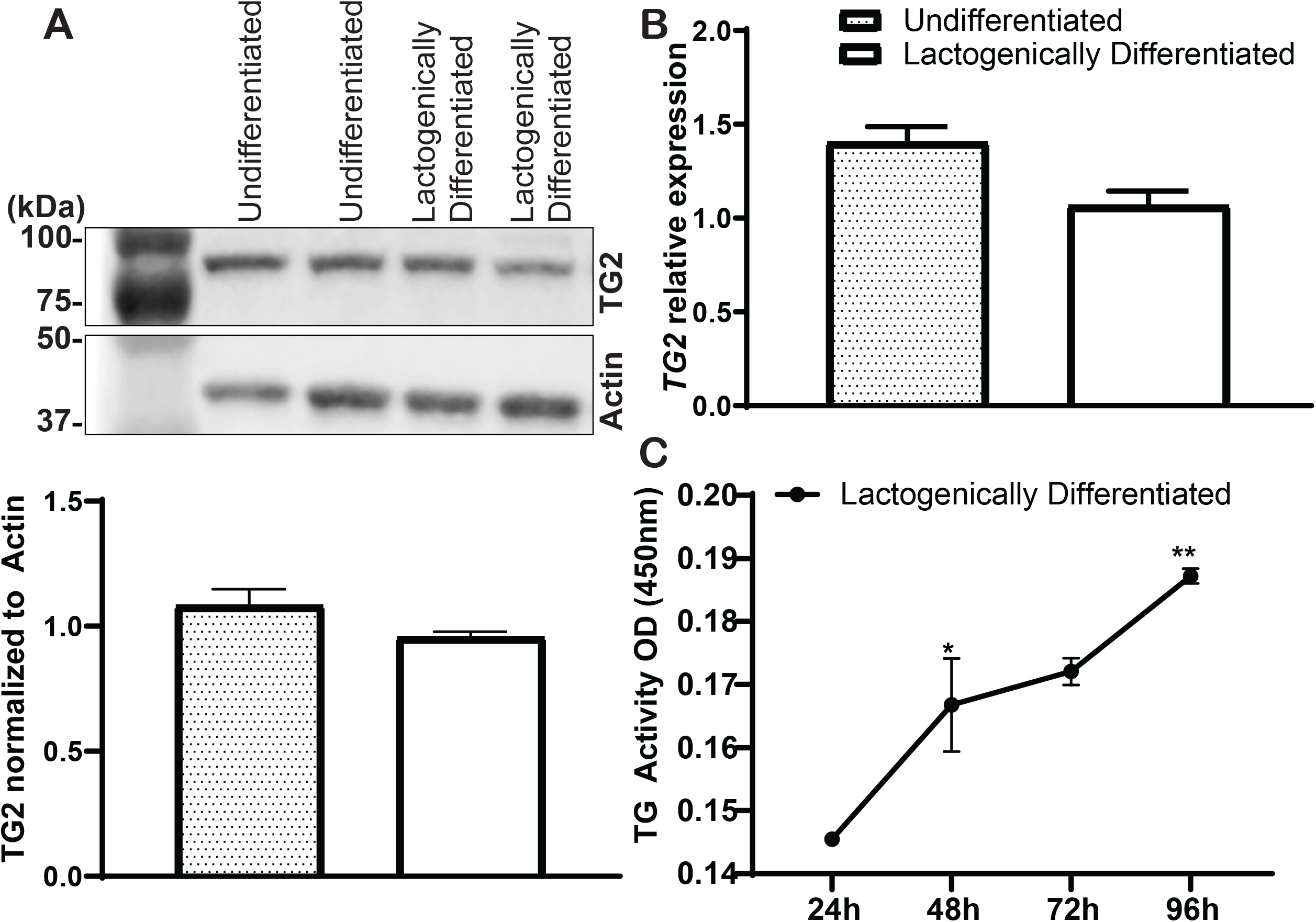
Transglutaminase protein and activity are observed in mouse mammary epithelial cells (HC11). **A.** Western blot analysis of transglutaminase 2 (TG2) in lysate of HC11 cells before differentiation (proliferation media) and after differentiation (lactogenic media) show no change in protein expression. TG2 quantification expressed relative to the actin control. **B.** *TG2* gene expression is unchanged before and after differentiation. **C.** TG activity increases overtime in the lactogenic differentiated cells: 72 hours (p=0.0194) and 96 hours (p=0.0022) compared to baseline.

### FLX and 5HTP increase intracellular 5HT concentrations resulting in increased synthesis of PTHrP

PTHrP (*Pthlh)* has been shown to be increased with increased 5HT. Fig. 2A depicts 5HT concentrations over time in lactogenic HC11 cells treated with FLX or 5HTP to increase intracellular 5HT. 5HT concentrations in 5HTP treated cells peak at 12 hours (Fig. 2B) whereas FLX treated cells have peak 5HT concentrations at 48 hours (Fig. 2C). We likely do not see an increase in intracellular 5HT in the 5HTP treated cells at 48 hours due to the degradation of 5HT by monoamine oxidase (MAO). We demonstrate a significant upregulation of *MAO-*A gene expression (Fig. 2D) in 5HTP treated cells, suggesting increased degradation of 5HT. We measured *SERT* (Fig. 2E) and *Tph1* (Fig. 2F) gene expression and saw increased expression of both genes when HC11 cells were treated with FLX, but 5HTP only increased *Tph1* mRNA expression. This confirms that only FLX treatment blocks SERT and both upregulates *de novo* synthesis of 5HT. At 48 hours post treatment, *Pthlh* mRNA expression (Fig. 2G) was significantly upregulated in FLX and 5HTP treated cells. Cyclic AMP (cAMP) is a common measure of PTHrP activity. At 48 hours post differentiation 5HTP significantly upregulated cAMP concentrations, whereas FLX tended to decrease cAMP concentrations, compared to lactogenic control (Fig. 2H). This suggests in the 5HTP treated cells *Pthlh* expression and activity is increased, whereas FLX treated cells only had increased *Pthlh* expression.

**Figure 2:**
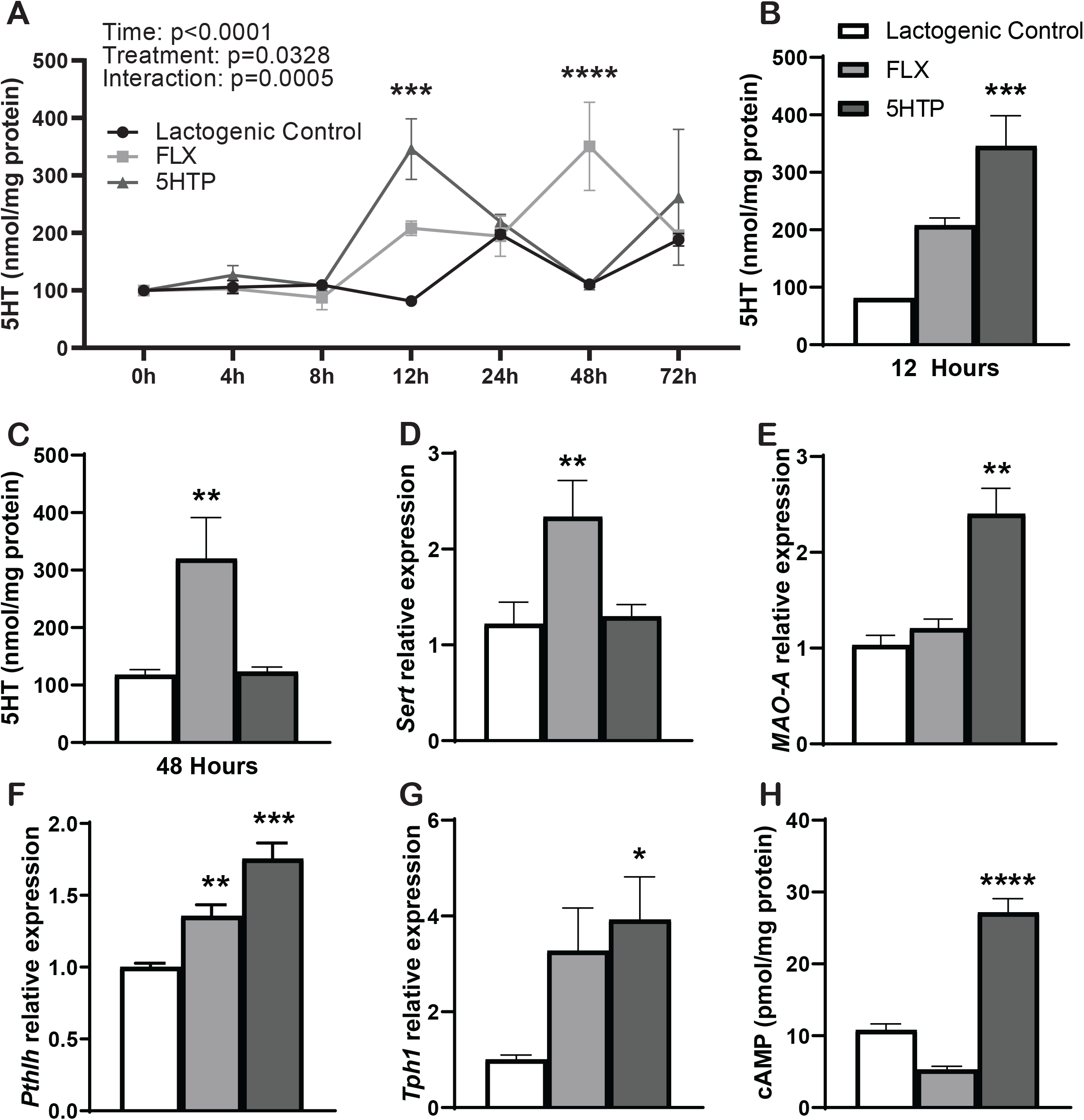
HC11 cells increase Pthlh gene expression and activity via cAMP with increases in intracellular serotonin (5HT) concentration. **A-C.** Intracellular 5HT concentration found a significant increase with time (p<0.0001), with treatment (p=0.0328), with an interaction between time and treatment (p=0.0002). 5HTP significantly increases at 12 hours post treatment (p=0.0005) compared to baseline and significantly increases (p=0.0017) compared to control at 12 hours **(B).** FLX reaches max 5HT concentration at 48 hours post treatment (p<0.0001) compared to baseline and is significantly increased (p<0.0001) compared to control at 48 hours **(C).** MDC does not impact 5HT concentration (not shown). **D.** Monoamine oxidase-A (MAO-A) gene expression is significantly upregulated in 5HTP treatment (p=0.0060) and not changed in FLX (p=0.6745). **E.** *Sert* gene expression in HC11 cells is upregulated with FLX treatment (p=0.0043) and unchanged in 5HTP (p=0.9988) **F.** *Tph1* gene expression is upregulated with FLX and 5HTP treatment (p=0.0702 and p=0.0191, respectively). **G.** *Pthlh* mRNA expression is significantly upregulated with FLX (p<0.0001) and 5HTP (p<0.0001) treatment. **H.** cAMP concentration significantly increases in 5HTP treated cells (p<0.0001) and FLX treated cells have decreased cAMP concentration (p=0.0864) at 48 hours compared to control.

### Inhibition of TG2 restores Pthlh gene expression to control levels in lactating HC11 cells

We examined the role of TG2 on induction of PTHrP in lactogenic mammary epithelial cells using a small molecule inhibitor, dansylcadaverine [MDC], to decrease serotonylation. Lactating HC11 cells had significantly upregulated *Pthlh* mRNA expression when treated with FLX or 5HTP and this was significantly decreased with TG inhibition using MDC (Fig. 3A). This result suggests TG2 is involved in 5HT’s regulation of *Pthlh*. Furthermore, we analyzed cAMP concentrations over time and observed significant changes across time, due to treatment, and the presence of interaction between time and treatment. Lactating HC11 cells treated with 5HTP exhibited increased intracellular cAMP concentrations at 24-, 48-, and 72 hours post lactogenic differentiation and FLX treated cells significantly decreased cAMP concentrations at 48 hours, with a tendency of decreased concentrations at 72 hours, compared to 0 hour. The lactogenic control had significantly decreased cAMP concentrations by 72 hours. Furthermore, cAMP concentrations (Fig. 3B) were unchanged with TG inhibition. TG2 gene expression was unchanged with treatment (Fig. 3C). TG activity (Fig. 3D) significantly changed over time, with treatment, and there was presence of an interaction of time and treatment. 5HTP significantly increased TG activity at 24 hours and FLX significantly increased activity at 72 hours compared to the lactogenic control. The lactogenic control had significantly increased TG activity at 72- and 96 hours post treatment, FLX treatment had significantly increased TG activity at 72 hours post treatment, and 5HTP had significantly increased TG activity at 96 hours post treatment compared to the 24-hour timepoint. MDC treatment did not result in significant changes when cells were treated with FLX or 5HTP, however MDC treatment increased TG activity at 96 hours post treatment. MDC exerts its inhibition by acting as an alternative substrate for TG, allowing TG to use MDC as a substrate instead of 5HT. We therefore would not expect MDC treatment to decrease TG activity.

**Figure 3:**
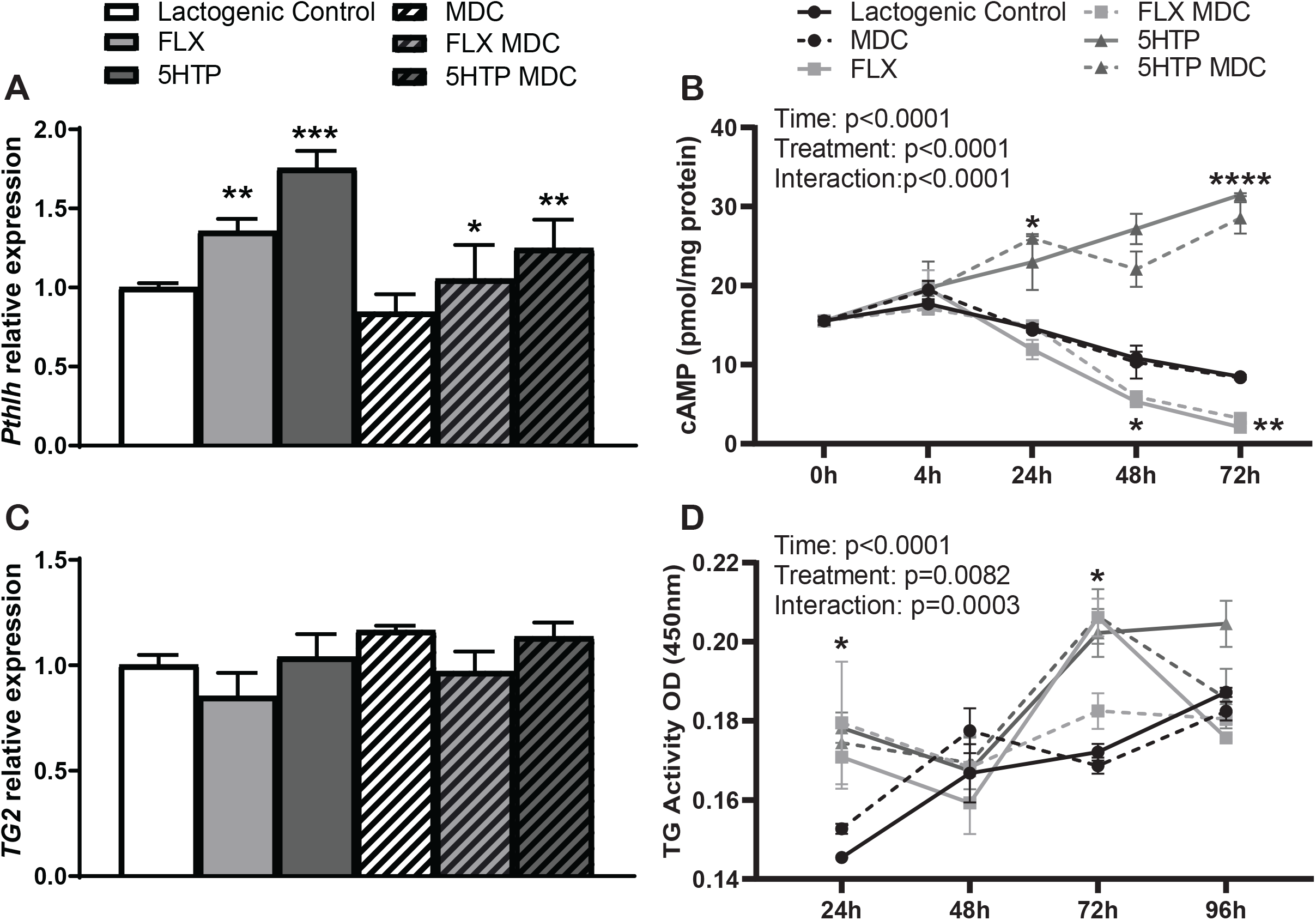
Inhibition of Transglutaminase restores *Pthlh* gene expression in HC11 cells. **A.** *Pthlh* mRNA expression is significantly increased in FLX (p=0.0050) and 5HTP (p=0.0006) compared to controls and is significantly decreased in FLX or 5HTP+ transglutaminase inhibition (MDC) (p=0.0400, p=0.0085 respectively) restoring *Pthlh* mRNA to lactogenic control levels. **B.** cAMP concentration is unchanged with transglutaminase inhibition. cAMP has a significant change over time (p<0.0001), treatment (p<0.0001), and interaction between time and treatment (p<0.001). 5HTP significantly increases at 24- hour, 48 hour, and 72 hours (p=0.0371, p=0.0029, p=0.0003 respectively) compared to baseline. 5HTP is significantly increased at 48- and 72 hour (p=0.0265, p<0.0001 respectively) and FLX is decreased at 48- and 72 hours (p=0.0176, p=0.0032 respectively) compared to lactogenic control. **C.** *Transglutaminase* mRNA expression is not changed with treatment. **D.** Transglutaminase activity is significantly changed with time (p<0.0001), treatment (p=0.0082), and the interaction between time and treatment (p=0.0003). Lactogenic control increases TG activity between baseline and 96 hours (p=0.0022), FLX increases TG activity between baseline and 72 hours (p=0.0350), and 5HTP increases TG activity between baseline and 96 hours (p=0.0166). MDC treatment kept TG activity constant with no significant changes. 5HTP was significantly increased at 24 hours (p=0.0461) compared to lactogenic control and FLX was significantly increased at 72 hours (p=0.0409) compared to lactogenic control.

### RhoA gene expression is upregulated when intracellular 5HT is increased and RhoA activity degrades over time

We assessed RhoA, Rab4a/b, and Rac1 as potential serotonylation target proteins in lactating HC11 cells. *RhoA* mRNA expression (Fig. 4A) was significantly upregulated with FLX and 5HTP treatments. FLX treatment in combination with the TG-inhibitor, MDC, restored *RhoA* mRNA to control levels, but 5HTP in combination with MDC did not alter *RhoA* mRNA expression. *Rab4a* (Fig. 4B) and *Rab4b* (Fig. 4C) expression were not altered by treatment. Next, we examined RhoA activity (Fig. 4D). RhoA activity was significantly affected by time and there was an interaction between time and treatment. RhoA activity was consistently decreased (p<0.0001) at 24-, 48-, and 72 hours compared to baseline for all treatments, likely due to degradation of Rho protein; FLX significantly decreased RhoA activity compared to control at 4 hours, but FLX and 5HTP treatments did not alter RhoA activity compared to control at any other time points. Due to the position of 5HT binding to the switch 2 domain on RhoA producing an inability to hydrolyze the bound GTP, it is possible that RhoA is constitutively active until it is degraded by the proteasome (41). Rac1 activity (Fig. 4D) was also affected by time and the interaction between time and treatment. Rac1 activity was significantly decreased but was not affected by treatment at any timepoint. Rac1 activity was decreased at 4 hours compared to 0 hours and plateaued at 4 hours and did not decrease due to time or treatment. Rac1 activity began to decrease at 4 hours, but plateaued, suggesting Rac1 is unlikely a serotonylation target.

**Figure 4:**
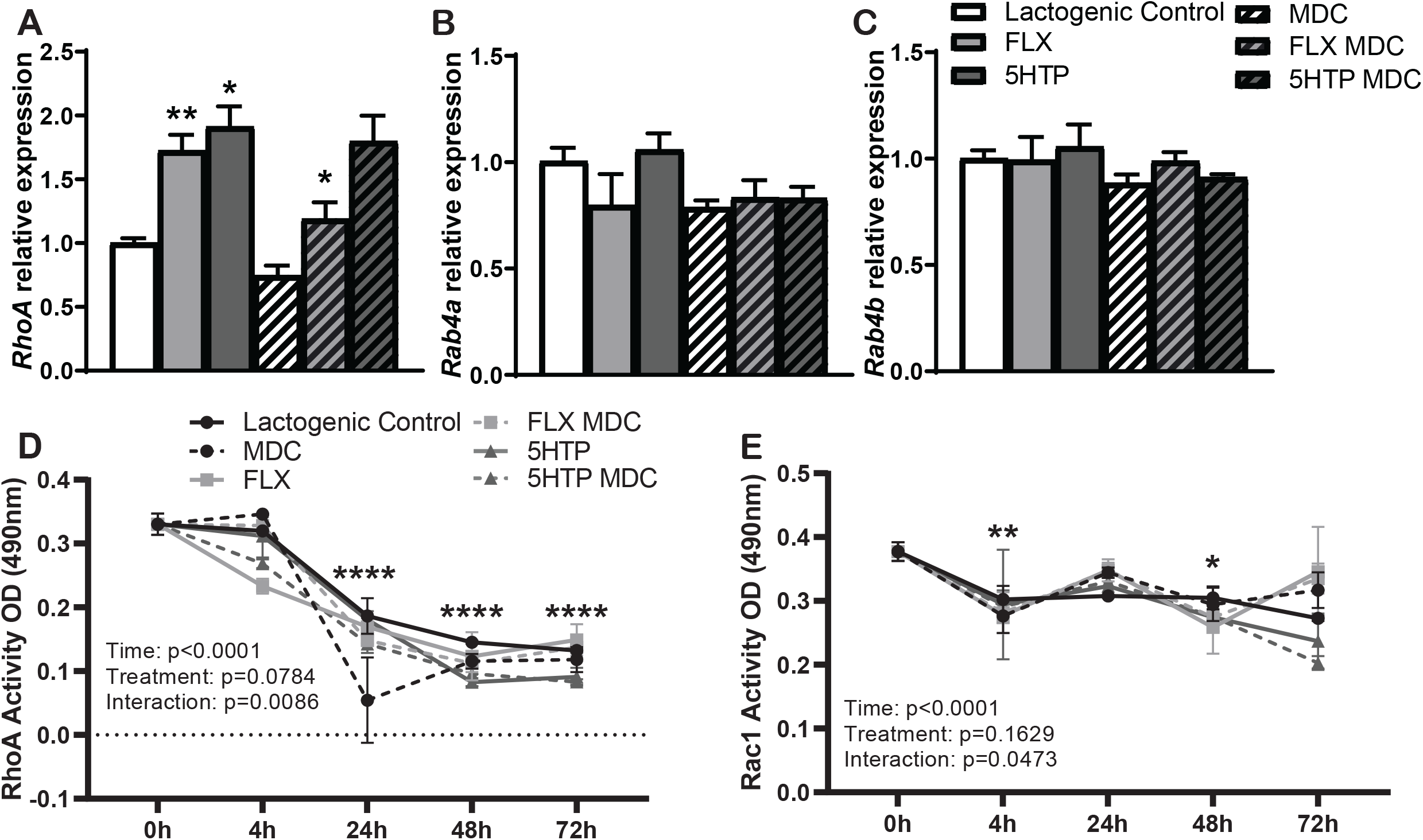
RhoA is serotonylated in HC11 cells. RhoA gene expression and activity are increased with increased intracellular 5HT. **A.** *Rho* mRNA expression is significantly upregulated with FLX (p=0.0022) and 5HTP (p=0.0100) compared to controls and is significantly decreased in FLX + MDC (p=0.0453). **B.** Rab4a mRNA expression remains unchanged with treatment. **C.** Rab4b mRNA expression remains unchanged with treatment. **D.** Rho activity decreases over time (p<0.0001), tendency of variation with treatment (p=0.0784), and a significant interaction between time and treatment (p=0.0086). FLX significant decreases activity at 4 hours (p=0.0275) compared to lactogenic control with no other changes with treatment. FLX significantly decreases activity at 4 hours (p=0.0087) compared to baseline. All treatments reduce activity at 24-, 48-, and 72 hours (p<0.0001 for all treatments and all timepoints) compared to baseline. **E.** Rac1 activity significantly decreases over time (p<0.0001), with no change with treatment (p=0.1629), and significant interaction between time and treatment (p=0.0473). There was no change with treatment compared to control and no change with MDC treatment. Control was significantly reduced at 72 hours (p=0.0082) compared to baseline, MDC reduced activity only at 4 hours (p=0.0116), FLX reduced activity at 4 hours (p=0.0126) and 48 hours (p=0.0023), FLX with MDC reduced activity at 4 hours (p=0.0179) and 48 hours (p=0.0096), 5HTP reduced activity at 48 hours (p=0.0098) and 72 hours (p=0.0003), and 5HTP with MDC reduced activity at 4 hours (p=0.0395) 48 hours (0.0102) and 72 hours (p<0.0001) compared to baseline.

## Discussion

Lactogenically differentiated mammary epithelial cells have been shown here and in previous studies to increase PTHrP, and further been demonstrated to be stimulated by increased 5HT (8,20,27). 5HT’s regulation of PTHrP is not fully understood, though mechanisms have been suggested such as action of 5HT through the 5HT2b receptor and altered DNA methylation of sonic hedgehog [SHH]; furthermore, during lactation there are interactions between PTHrP and the calcium sensing receptor (8,27,42,43). In this study we examined the molecular process of serotonylation as a potential mediator of increased PTHrP. To our knowledge, this is the first study examining serotonylation in normal mammary epithelial cells that have been differentiated with lactogenic hormones. Our data suggest a novel potential mechanism of 5HT’s regulation of PTHrP is serotonylation.

We utilized a commonly used mouse mammary epithelial cell line (HC11) to study *in vitro* lactogenesis. Upon stimulation with lactogenic hormones (prolactin, insulin, and hydrocortisone), these cells undergo lactogenic differentiation characterized by the formation of mammospheres and upregulation of milk protein genes, (supplemental Fig. 1) (44–46). We demonstrated the presence of protein and gene expression of TG2 in HC11 cells but observed that they are unchanged due to stimulation with lactogenic hormones. This is consistent with studies implicating TG2 in mammary epithelial breast cancer (40,47). Further we characterized TG activity, finding that as the differentiation of the mammary epithelial cells increase, the activity of TG increases. TG2 is a calcium dependent enzyme; therefore, due to the demand of calcium by the mammary gland for milk synthesis, it is possible that the influx of calcium is potentially allowing TG to have increased activity during lactation (13,30).

We further went on to manipulate 5HT concentrations using two pharmacological mechanisms: treatment with FLX or 5HTP. We chose FLX and 5HTP rather than 5HT due to the short half-life and rapid degradation of 5HT, and the use of 5HTP or FLX have been utilized by our lab to increase 5HT in many previous studies (10,20,48–50). We demonstrated that intracellular 5HT concentrations increase over time with either 5HTP or FLX treatment, with peak 5HT concentrations being achieved at 12 or 48 hours, respectively. At 48 hours, when these cells have fully differentiated, we no longer see an increase in 5HT concentration with 5HTP treatment, likely due to the degradation of 5HT by monoamine oxidases [MAO], as measured by a significant increase in *MAO-*A gene expression with 5HTP treatment (51,52). As expected, this increase in intracellular 5HT concentration with either 5HTP or FLX treatment sufficiently upregulated PTHrP mRNA expression, consistent with previous data (8,20,49). Consistent with our hypothesis, when we combined 5HTP or FLX treatment with a TG-inhibitor, MDC, PTHrP expression was decreased. This suggests TG2-dependent serotonylation is one mechanism likely involved in 5HT’s regulation of PTHrP synthesis. Many have attributed MDC’s inhibitory effects on signaling pathways to the process of serotonylation (33,35,53).

PTHrP activity has commonly been measured using cAMP concentration which is downstream of the PTH-receptor (20,54,55). We found 5HTP significantly increased cAMP concentrations at 24-, 48-, and 72- hours post treatment compared to the lactogenic control. However, FLX significantly decreased cAMP concentration at 48 hours, and had a tendency to decrease cAMP concentrations at 72 hours compared to lactogenic control. This is in contrast to our previous results *in vivo* FLX exposure increased mammary gland cAMP concentrations on day 10 of lactation, after approximately 14 days of treatment with FLX (20). It is possible that duration and timing of treatment (acute vs. chronic) with these compounds can result in different effects on the mammary gland or other tissues, which we and other have previously shown (48,56). Interestingly, in the human breast cancer cell line, MDA-MB-231, a similar observation was found regarding decrease in cAMP with an increase in 5HT and a switch from growth inhibition to stimulation was attributed to the altered cAMP dynamics (57). MDC treatment had no significant effects PTHrP activity when measuring cAMP concentrations.

MDC did not completely abolish PTHrP expression or reduce cAMP concentrations at the low concentration used in this study (25μM), which is substantially lower than many studies which range from 200μM up to 5mM concentrations (36,41,53,58). We chose the lower concentration due to the chronic duration of treatment (48+ hours) compared to acute duration (1-8 hours) and studies have shown effects at lower concentrations. One study observed that 20μM MDC treatment overnight resulted in a significant decrease in 5HT mitogenesis of distal primary bovine arterial smooth muscle cells, though higher doses (up to 200μM) resulted in a further decrease (53). Another study found 25μM MDC treatment for 3 hours in L6 rat muscle cells results in significant decrease in 5HT induced effects on GLUT4 translocation, glucose uptake, and glycogen content (33). We therefore concluded a lower dose would sufficiently inhibit serotonylation, but it is possible that we would see further reductions in PTHrP or cAMP if we used a higher concentration of MDC.

We then characterized TG activity, finding it increases over time and is altered with treatment at various timepoints (5HTP at 24 hours and FLX at 72 hours compared to lactogenic control). TG is implicated in many other cellular processes, including monoaminylation (e.g. serotonylation, histaminylation, dopaminylation, and norepinephrinylation) and protein-protein crosslinking involved in extracellular matrix proteins, growth factor activity, integrin activity, oxidative stress/inflammation, and EGF/EGFR signaling in epithelial cancer cells (59–62). It is possible that by increasing 5HT concentrations we are increasing serotonylation through increased substrate (5HT) availability. As expected, treatment with MDC resulted in no change in TG2 activity. MDC is a small-molecule inhibitor that acts as an alternative substrate exploiting the protein-crosslinking activity of TG (63). It has been shown that MDC will outcompete the donor substrate (5HT), thereby reducing serotonylation while keeping TG activity steady (63,64).

G-proteins have emerged as the most common serotonylation target. Herein, we probed a variety of G-proteins including: RhoA, Rab4a/b, and Rac1. We have determined RhoA is the most likely serotonylation target in lactating mammary epithelial cells, though it is unclear whether serotonylation of RhoA is involved in the regulation of PTHrP. We began by examining gene expression of *RhoA, Rab4a,* and *Rab4b* and determined that *RhoA* as the only gene that exhibited changes in expression. This increase in transcription of RhoA may be a mechanism to increase Rho protein in response to the degradation of serotonylated Rho protein. A previous serotonylation study in pulmonary hypertension in smooth muscle cells have shown Rho protein significantly decreases at 24-, 48-, and 72- hours post 5HT stimulation, attributed to the degradation to eliminate activity (41). When G-proteins, such as the Rho family and Rab family, are serotonylated, they lose the intrinsic ability to hydrolyze GTP to GDP, thus rendering them constitutively active until proteasomal degradation (29,30,37,65). Consistent with what was previously described regarding serotonylated Rho, when examining RhoA activity, we found a consistent, significant decrease at 24-, 48-, and 72- hours, Rac1 activity, on the other hand, remained relatively constant from 4- to 48 hours, with only a significant decrease from baseline at 4 hours and 72 hours. This suggests Rac1 undergoes minimal degradation, suggesting it is unlikely to be serotonylated. Although we do not see a change in RhoA activity with treatment, the overall decrease in Rho activity coupled with the increase in gene transcription, suggests Rho may degraded to inactivate it and transcription increases to replenish the degraded Rho protein. We have not confirmed Rho’s involvement in PTHrP synthesis to date.

To our knowledge, this study is the examining serotonylation in normal mammary epithelial cells under lactogenic conditions. We conclude that 5HT regulates PTHrP in part through the novel mechanism of serotonylation. PTHrP synthesis is tightly regulated through many mechanisms in the mammary epithelial cells including 5HT receptor 2b [5HT2b] activation, altered DNA methylation of sonic hedgehog [SHH], calcium sensing receptor, and now serotonylation.

## Methods

### Cell Culture

Mouse mammary epithelial cells (HC11), a prolactin responsive cell line that undergoes lactogenic differentiation were utilized for this experiment. HC11 cells were plated at a seeding density of 250,000 cells/well of a 12 well plate or 1,000,000 cells/10cm plate, sustained in proliferation media (RPMI 1640, 10% FBS, 1% antibiotics, 5μg/mL insulin, 25ng/mL EGF), 48 hours later the cells were confluent. Once confluent, EGF was removed (proliferation media: RPMI 1640, 10% FBS, 1% antibiotics, 5μg/mL insulin) for 48 hours to initiate the lactogenic differentiation. After 48 hours, lactogenic media (RPMI 1640, 10% FBS, 1% antibiotics, 10μg/mL insulin, 1μg/mL prolactin, 0.5μg/mL hydrocortisone) was added to induce lactogenic differentiation. All treatments are as following: 40μM fluoxetine (Sigma Aldrich, catalog number F132, St. Louis, MO), 500 μM 5HTP (Sigma Aldrich, catalog number H9772, St. Louis, MO), 25μM dansylcadaverine dissolved in DMSO (Sigma Aldrich, catalog number 30432, St. Louis, MO), or DMSO control was added to the lactogenic media and after 48 hours the cells were harvested. Cell culture differentiations were done in duplicate and this was repeated three times.

### RNA and RTqPCR

HC11 cells were harvested with TRI-Reagent (Molecular Research, Thermo Fisher Scientific, catalog number NC9277980, Waltham, MA) according to manufacturer’s protocol. 1μg RNA was reverse transcribed with High-Capacity cDNA Reverse Transcription Kit (Applied Biosystems, Thermo Fisher Scientific, catalog number 4368814, Waltham, MA) with murine RNase inhibitor (New England Biolabs, catalog number M0314L, Ipswich, MA). Quantitative RT-PCR was conducted using the CFX96 Touch-Real-Time PCR Detection System (Bio-Rad Laboratories, Hercules, CA). Reaction mixtures and cycling conditions were performed as previously described (49). Primers were designed with an optimal annealing temperature of 60°C. Amplification efficiencies of primers were accepted within a range of 95-105% and a singular melt-curve. The primer sequences are listed in Table 1. The housekeeping parameter was the geometric mean of *Rsp9* and *S15*. Analysis was conducted using the 2^−ΔΔCT^ method.

**Table 1.**
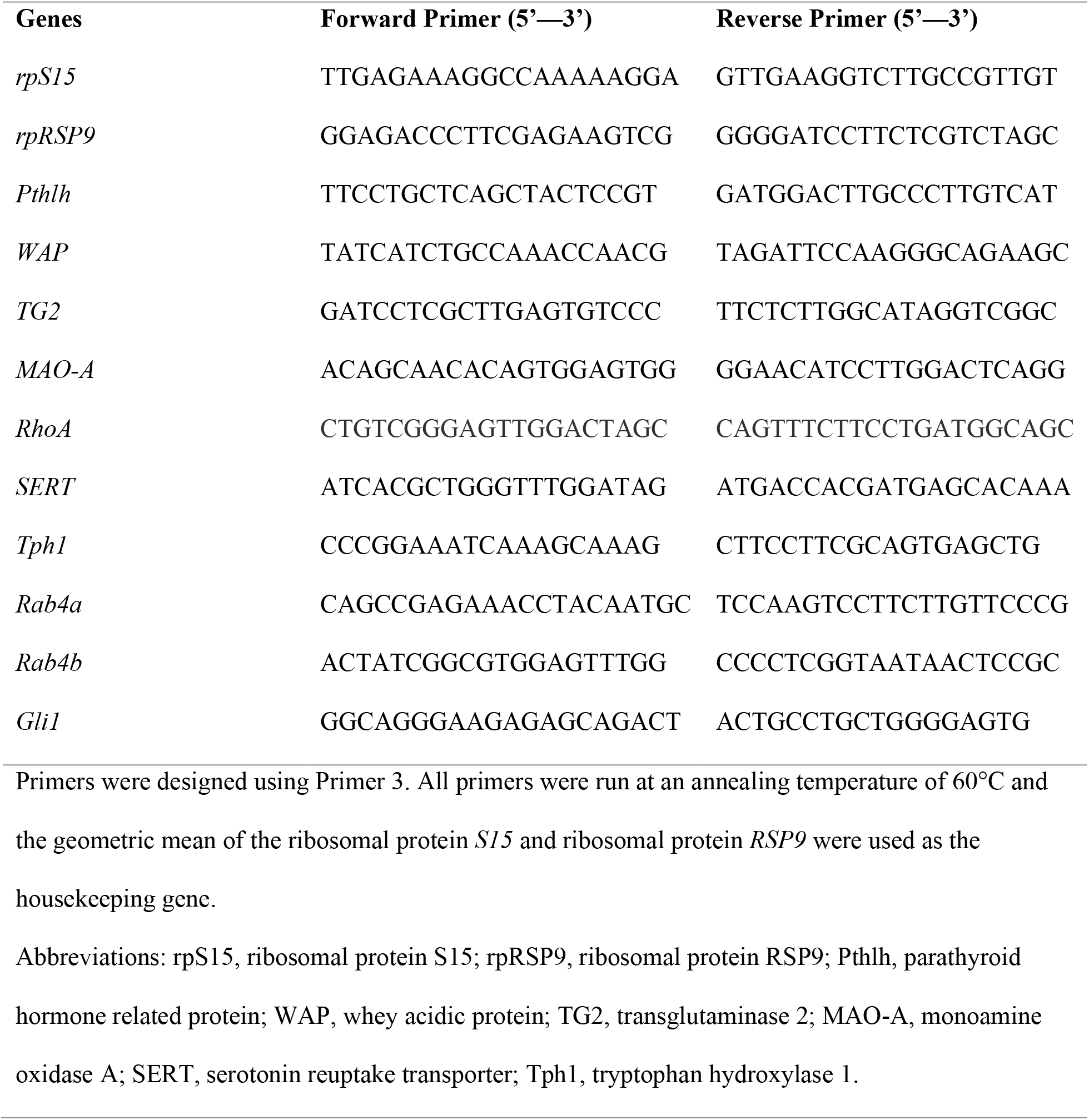
Primer Sequences for the studied genes quantified by real-time-PCR

### Protein Extraction and Immunoblotting

HC11 cells were harvested with radioimmunoprecipitation assay buffer (1X PBS, 1% nonidet P-40, 0.5% sodium deoxycholate, 0.1% SDS) supplemented with 10μL/mL Halt Protease and Phosphatase Inhibitor Cocktail (Thermo Fisher Scientific, catalog number 78441, Waltham, MA). Lysates were homogenized and cleared with centrifugation for 15 minutes at 12,000xG. Protein concentration was determined using bicinchoninic acid assay (Bioworld, catalog number 20831001-1, Dublin, OH). Protein lysates were diluted to 1.5μg/μl with 5x sample buffer containing SDS and β-mercaptoethanol and was heated at 95°C for 10 minutes. 15μg protein was separated by electrophoresis on a gradient (8-20%) SDS-polyacrylamide gel and transferred for 1 hour at 100V onto a polyvinylidene difluoride membrane (Millipore Sigma, catalog number IPVH00010, Burlington, MA). Membranes were blocked for 1 hour with Sea Block Blocking Solution (Thermo Fisher Scientific, Waltham, MA) and probed overnight at 4°C with 1:1000 TG2 (Abcam, catalog number ab421, Cambridge, United Kingdom) and 1:1000 β-actin (Cell Signaling Technology, catalog number 4967S, Danvers, MA). The following day, the membrane was washed 3 times with TBST and probed 1:5000 with fluorescent secondary antibodies (Li-Cor Biosciences, IRDye 800 CW catalog number 925-32213, IRDye 680 RD catalog number 925-68070, Lincoln, NE), then washed 3 times with TBST. Protein bands were detected using Li-Cor Odyssey Fc (Li-Cor, Lincoln, NE) with a 2-minute exposure for 700 channel and 10-minute exposure for 800 channels. Image analysis and protein band quantification were performed using Image Studio Lite software (Li-Cor Biosciences, version 5.2, Lincoln, NE).

### Assays

Intracellular 5HT concentrations were determined using a Serotonin Enzyme Immunoassay Kit (Beckman Coulter, catalog number IM1749, Brea, CA) using 50μg total protein per the manufacturer’s instructions. Intracellular cAMP concentrations were determined using a cyclic AMP XP Assay Kit (Cell Signaling Technology, catalog number 4339S, Danvers, MA) using 50μg total protein per the manufacturer’s instructions. Intracellular transglutaminase activity was determined using a Transglutaminase Assay Kit (Sigma-Aldrich, catalog number CS1070, St. Louis, MO) using 20μg total protein per the manufacturer’s instructions. Intracellular RhoA activity was determined using a RhoA G-LISA Activation Assay Kit (Colorimetric format) (Cytoskeleton, catalog BK124, Denver, CO) using 20μg total protein per the manufacturer’s instructions. Intracellular Rac1 activity was determined using a Rac1 G-LISA Activation Assay Kit (Colorimetric format) (Cytoskeleton, catalog BK128, Denver, CO) using 25μg total protein per the manufacturer’s instructions. All assays had an intraassay CV of <10% and the interassay CV of <5%.

### Statistical Analysis

All statistical analyses were conducted using Graph Pad Prism 8 (Version 8.4.0). Analysis between two treatments were performed using a Student’s unpaired two-sided *t* test and analysis between multiple treatments were performed using one-way ANOVA. Bartlett’s test was applied to test equal variances among treatments. If sample populations did not have equal variances, Welch correction for unequal variance was applied. Analyses with multiple time points were conducted using a two-way ANOVA with repeated measures with Tukey’s multiple comparisons test to detect differences between treatment groups. For all analyses, differences between the mean were considered significant if P<0.05. All values are reported as means +/− SEM.

## Data Availability

All data are contained within the manuscript.

## Abbreviations used are

5HT: serotonin
TPH: tryptophan hydroxylase
5HTP: 5-hydroxytryptophan
PTHrP: parathyroid hormone related protein
SSRI: selective serotonin reuptake transporter
FLX: fluoxetine
TG: transglutaminase
MDC: (mono)dansylcadaverine
MAO: monoamine oxidase
SERT: serotonin reuptake transporter
cAMP: cyclic adenosine monophosphate;

## Acknowledgements

We thank Luma C. Sartori for her technical assistance. We thank Linda S. Schuler for providing us with the HC11 cells.

## Supplemental Figure

**Supplemental Figure 1.**
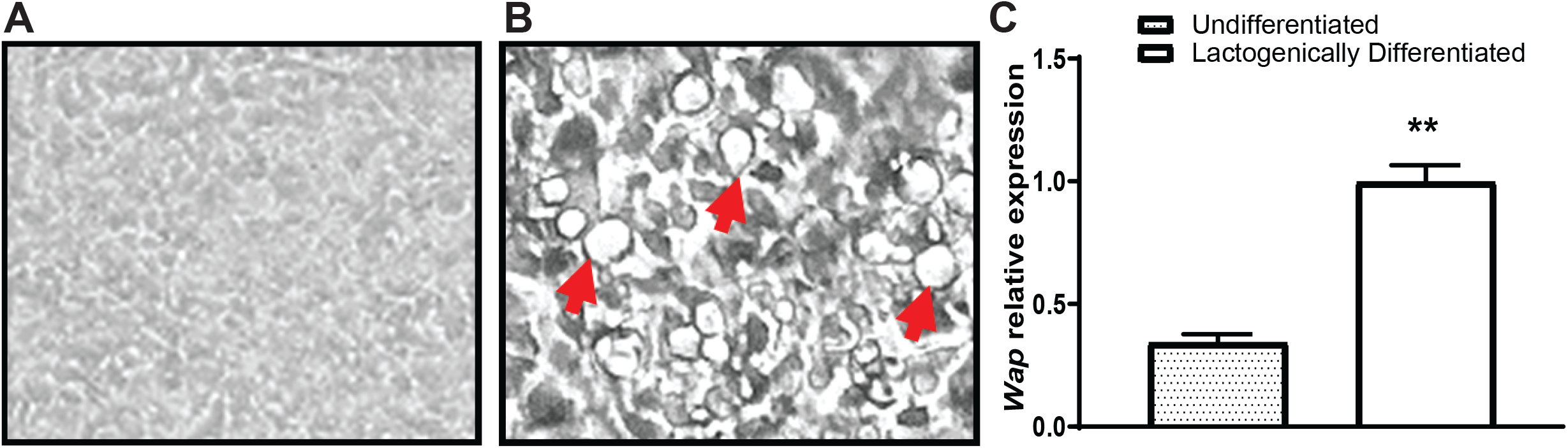
HC11 cells undergo lactogenic differentiation when treated with lactogenic hormones (prolactin, hydrocortisone, and insulin). Undifferentiated HC11 cells **(A)** are an epithelial monolayer, whereas during lactogenic differentiation **(B)** the cells undergo a morphological change, including formation of mammospheres (arrows). Differentiated cells begin to synthesize milk protein genes; whey acidic protein [WAP] **(C)** a milk protein gene, is significantly increased (p=0.004) in lactogenic differentiated cells compared to undifferentiated.

